# Transcriptome analysis of the quantitative distribution of the *Cyrtotrachelus buqueti* population in two cities in China

**DOI:** 10.1101/153148

**Authors:** Chaobing Luo, Anxuan Liu, Wencong Long, Hong Liao, Yaojun Yang

## Abstract

**Background:** *Cyrtotrachelus buqueti* is a forest pest that severely damages bamboo shoots. Reducing the population of this insect involves complex mechanisms and is dependent on diverse gene expression influenced by environmental factors.

**Methods:** In this study, samples from two regions of China, Muchuan in Sichuan Province and Chishui in Guizhou Province, were investigated through RNA-seq to explore the causes and molecular mechanisms underlying the population reduction of this species. Environmental factors, such as temperature, heavy metal content, and pH, may affect the reduced population of *C. buqueti* in Chishui.

**Results:** Approximately 44 million high-quality reads were generated, and 94.2% of the data were mapped to the transcriptome. A total of 15,641 out of the 29,406 identified genes were predicted. Moreover, 348 genes were differentially expressed between the two groups of imagoes and included 77 upregulated and 271 downregulated UniGenes. The functional analysis showed that these genes were significantly enriched in ribosome and metabolic pathway categories. The candidate genes, which contributed to *C. Buqueti* reduction, included 41 genes involved in the ribosome constitution category, five genes in the one-carbon pool pathway by folate category, and five heat shock protein genes.

**Conclusions:** Downregulation of these candidate genes seems to have impaired metabolic processes, such as protein, DNA, RNA, and purine synthesis, as well as carbon and folate metabolism, and finally resulted in the observed reduced population of *C. buqueti*. Furthermore, temperature, heavy metal content, and pH might influence the population by altering the expressions of genes involved in these metabolic processes.

## 1. Introduction

*Cyrtotrachelus buqueti* belongs to the class of insects in the order Coleopterai, the family Curculionoidea, and the genus *Cyrtotrachelus*. It is found in southern China and several countries of Southeast Asia, such as the Socialist Republic of Vietnam, Thailand, and the Union of Myanmar (Rui-Ting, 2005). *C. buqueti* is a pest in cluster bamboo forests and seriously limits the development of bamboo products, adversely affecting the income of bamboo farmers. This species damages bamboo shoots; specifically, the larvae bore into cluster bamboo shoots, such as *Phyllostachys pubescens*, *Neosinocalamus affinis*, *Bambusa textilis*, and *Dendrocalamus farinosus*. For this reason, the insect has been listed as a dangerous forestry pest by the State Forestry Administration of China since 2003 (Yang et al., 2009).

*C. buqueti* produces one generation per year in Sichuan and overwinters inside puparia in the soil. The adult emerges from the soil from June to October, which is almost the same time period as bamboo shooting. The activity period of adults lasts for around one month. The adult starts to mate and oviposit after 2 days of feeding. Female adults lay eggs in bamboo shoots, using their long, hard mouths. The larva causes damage to bamboo shoots from July to September and pupates sometime between the last 10 days of July to the last 10 days of October (Wang et al., 2005, Nie, 2010). The high content of special volatiles, primary benzaldehyde, and the special proportion of components on the top of the tufting bamboo shoot could produce an important odor signal that attracts *C. buqueti* toward the plant (Yang et al., 2010). Feeding and oviposition behaviors enhance the difficulty forecasting population density and injury percentage of bamboo shoots. There are various methods to control the *C. buqueti,* for example, sapling quarantine, removing the puparia from the soil and the larva from the bamboo shoot manually, pesticides, and biological control, such as benzaldehyde (Nie, 2010). However, none of the methods are perfect, and are either inefficient or costly.

Based on our investigation, the *C. buqueti* hazard rate is more than 40% across the 67,000-hectare bamboo cluster forests at Muchuan and Mabian, Sichuan province. However, in Chishui, Guizhou province and its surrounding areas, such as Hejiang in the Sichuan province, only slight damage was observed and the hazard rate was less than 1% in 82,000-hectare bamboo clusters, which is less than 300 kilometers away from the first site (Yang, 2011). The reasons for and effects of this phenomenon are not fully understood. At present, many forecast studies have been launched on insects like *Drosophila* and mosquitoes (Guruprasad et al., 2010, Wegbreit and Reisen, 2000) in order to study their population change and biological damage driven by agriculture against the background of global environmental change (Clark et al., 2001). In addition, next generation high-throughput DNA sequencing techniques, such as Illumina sequencing, provide the opportunity for research in life science with considerable cost savings and efficiency (Ansorge, 2009). *C. buqueti* has been investigated in several studies, but the molecular regulation mechanisms are poorly understood and relevant genomic resources are scarce. Therefore, we sequenced and annotated the transcriptome of *C. buqueti* from the two study areas to screen for any genes that may be responsible for the reduction in the quantitative distribution of the insect, using Illumina HiSeq 2500 sequencing, with the goal of discovering safer prevention methods.

## 2. Materials and Methods

### 2.1 Insect material preparation

*C. buqueti* Guerin, from Muchuan (N103°98′, E28°96′) and Chishui (N105°69′, E28°57′), were used for transcriptome analysis. Six male adults were sampled. The digestive, reproductive, and musculature systems of *C. buqueti* were removed and mixed. The mixtures were transferred immediately to liquid nitrogen and stored subsequently at −80°C until RNA extraction. The digestive, reproductive, and musculature systems of *C. buqueti* were sampled from three comparable insects from same place using three biological replications. The Muchuan and Chishui samples were used to construct six libraries, which were named C2, C3, C4, D2, D3, and D4.

### 2.2 Detection of temperature, heavy metal concentration, and pH

The annual mean temperature, monthly mean maximum temperature, and monthly mean minimum temperature were derived from the TIANQI network (www.tianqi.com). Inductively coupled plasma mass spectrometry and potentiometry were performed to detect heavy metal concentration and pH, respectively.

### 2.3 RNA isolation, library construction, and sequencing

Total RNA was extracted using the RNAprep pure Tissue Kit (DP431, TianGen Biotechnology, Beijing, China) and was treated with RNase-free DNase I (TianGen Biotechnology, Beijing, China) to remove genomic DNA contamination. Briefly, 1.0% agarose gel stained with Gel-Red was evaluated to determine the RNA integrity. A K5500 spectrophotometer (KAIAO, Beijing, China) and an Agilent 2100 Bioanalyzer (Agilent Technologies, CA, USA) were used to assess RNA quantity and quality. The RNA integrity number (RIN) was greater than 8.0 for all samples (Clerico et al., 2015). RNA samples from the three systems for each group were pooled together in equal amounts, in order to generate one mixed sample. These six mixed RNA samples were used to construct the cDNA library and Illumina sequencing, which was managed by Beijing ANOROAD Bioinformatics Technology Co., Ltd. The oligo-dT beads were used to isolate mRNA, which were then fragmented into short sections and added into the fragmentation buffer. The short mRNA fragments were used as templates, with random hexamer primers, to synthesize first-strand cDNA. Next, dNTPs, DNA polymerase I, and response buffer were used to synthesize the second-strand cDNA. The double-stranded cDNAs were purified using the QIAQuick PCR kit, eluted with ethidium bromide, and then used for “A” base addition and end-reparation. The cDNAs were finally ligated with sequencing adapters and a 1.0% agarose gel stained with G was used to fragment recovery.

### 2.4 Sequence tag preprocessing and mapping

The sequence tag was preprocessed according to a previously described protocol (Li et al., 2009, Langmead et al., 2009). Raw reads were cleaned by removing reads with adaptors, low quality (≤19%), or Ns (>5%). Clean reads were mapped to transcript sequences after assembly with Bowtie 2 software, allowing for a maximum of 2 nucleotide mismatches (Tatusov et al., 1997).

### 2.5 Gene expression calculation and pathway analysis

Gene expression was calculated using RPKM (Mortazavi et al., 2008). The DEseq package (ver. 2.1.0) was used to detect DEGs between the two sample groups (Anders and Huber, 2010, Wang et al., 2010). DEGs were selected based on |log2 Ratio|≥1 and a Padj value < 0.05. FDR was used to determine the P-value threshold in multiple tests. The absolute value of the log 2 (fold change) with RPKM ≥ 1 and an FDR ≤0.005 were used as the thresholds in the study to determine significant differences in gene expression.

### 2.6 Functional analysis of DEGs

Functional enrichment analyses, such as GO and KEGG, were performed in order to identify whether DEGs were significantly enriched in GO terms or metabolic pathways. Blast2GO was performed to implement the GO enrichment analysis of DEGs. The corrected P value of GO terms (P< 0.05) was considered significantly enriched by DEGs. The statistical enrichment analyses of differential expression genes were calculated using KOBAS software for the KEGG pathways. The GENES database for KEGG contained the whole genome, whether complete or incomplete, the PATHWAY database included gene functional statistics, and the LIGAND Database contained the enzyme database. Pathways with an FDR value ≤0.05, which were defined as those with genes, showed significant levels of differential expression.

### 2.7 Quantitative real-time PCR validation of RNA-Seq data

Twenty DEGs were chosen for validation with quantitative real-time PCR (RT-qPCR). The primers, which were designed with the Primer 3.0 software (http://biotools.umassmed.edu/bioapps/primer3_www.cgi), are listed in Table 1. RT-qPCR reactions were analyzed in the ABI StepOneTM Plus Real-Time PCR System with SYBR Green PCR Master Mix (TianGen, Beijing, China), and amplified with 1μL of the cDNA template, 10μL 2×SupperReal PriMix Plus, 2μL of 50×ROX Reference Dye, 0.5μL of each primer (20 μmol/μL), and a final volume of 20μL was achieved by adding water. The amplification program consisted of one cycle of 95 °C for 15 min, followed by 30 cycles of 95 °C for 5 s, 60 °C for 20 s and 72 °C for 20 s. Fluorescent products were detected in the last step of each cycle. The melting curve analysis was performed at the end of 30 cycles, in order to ensure proper amplification of target fragments. All RT-qPCR for each gene was performed in three biological replicates, with three technical repeats per experiment. Relative gene expressions were normalized by comparison with the expression of lotus β-actin (c28453_g1_i3), and were analyzed using the 2^−ΔΔC^T method. The data are indicated as mean ±SE (n=9). Statistical analysis of the RT-qPCR data was conducted using the ANOVA procedure in SAS 8.1 (SAS Institute, Cary, NC, USA).

**Table 1.**
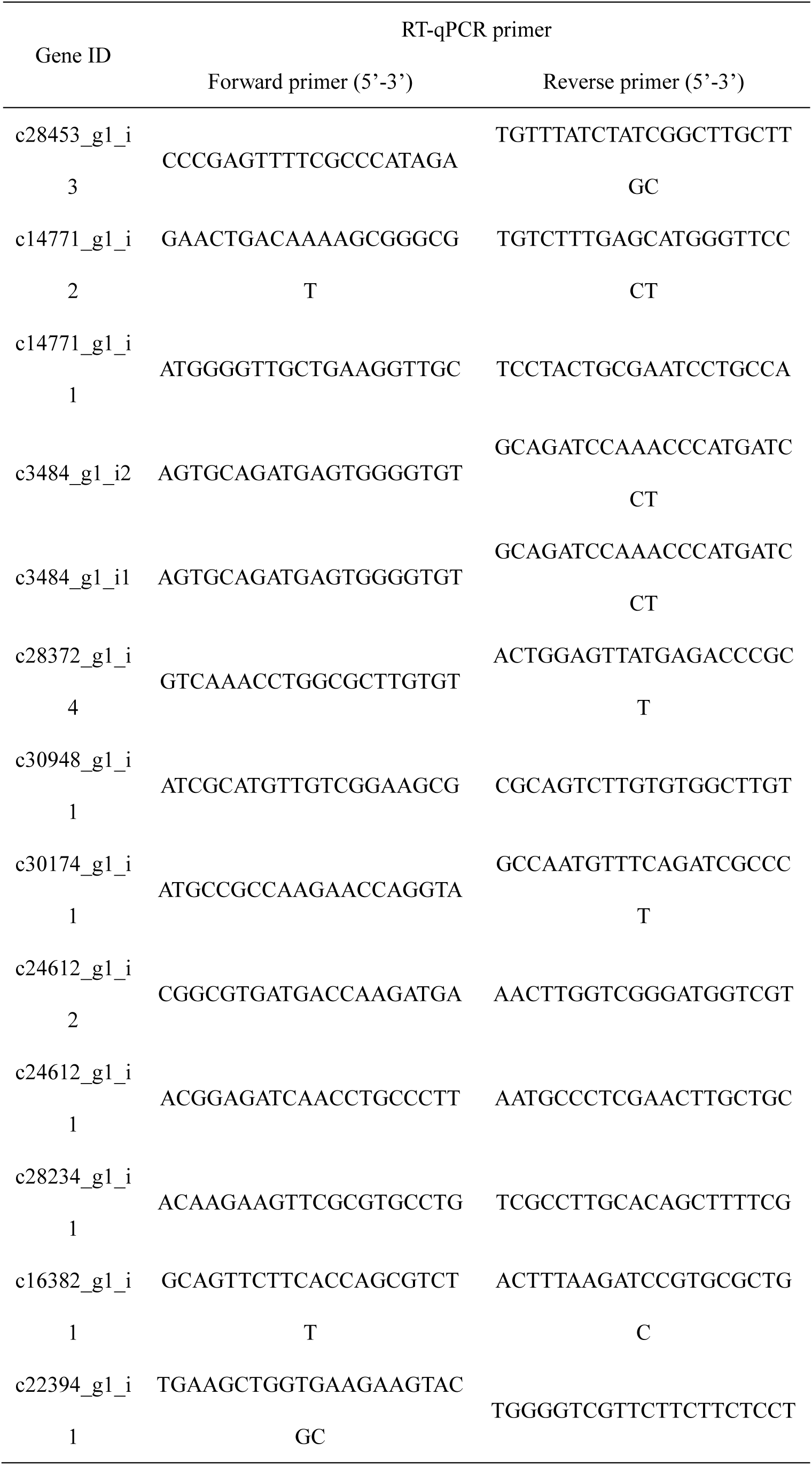
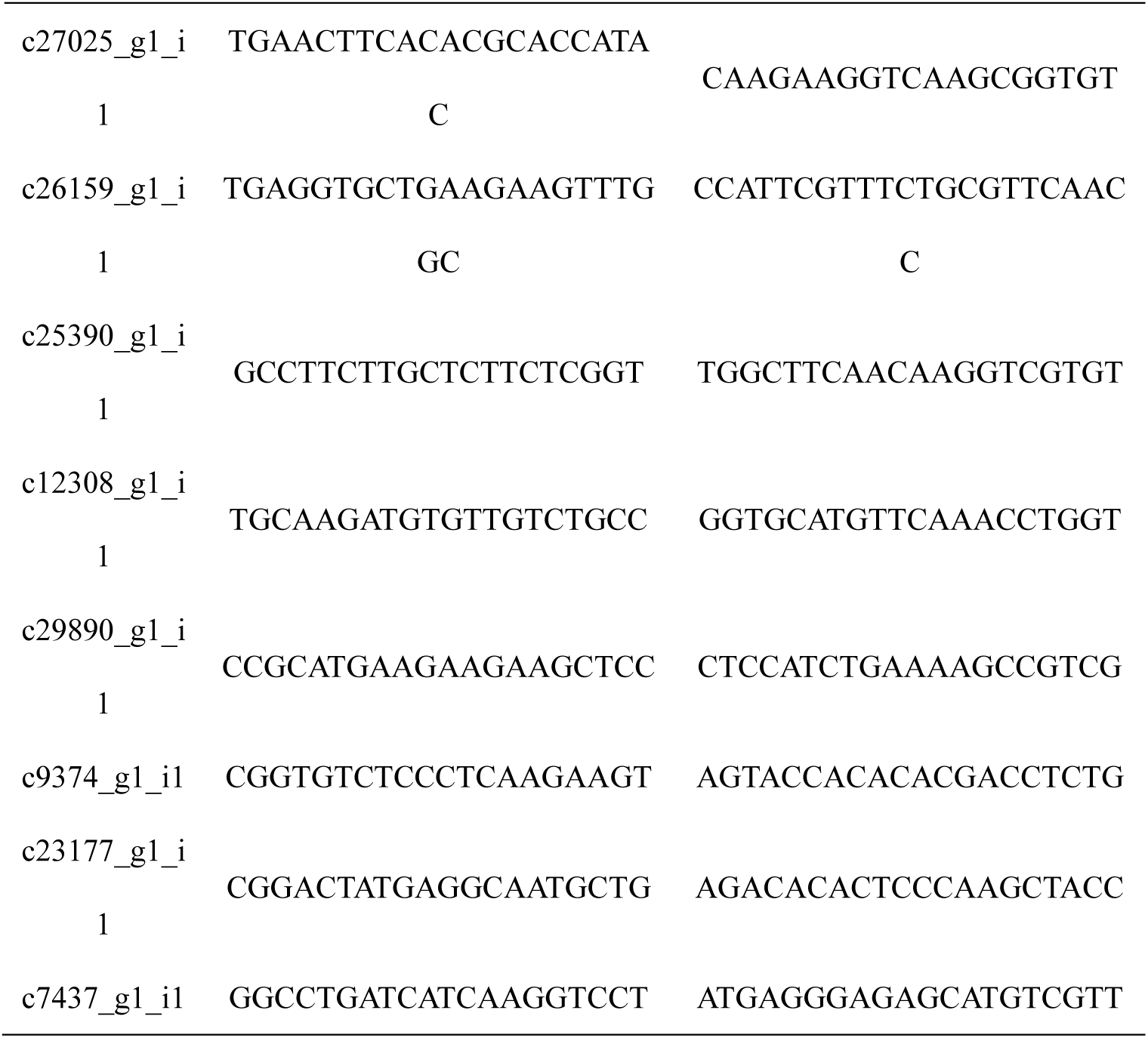
Primers used in RT-qPCR

## 3 Results

### 3.1 Mortality and living environment analysis of C. buqueti in Chishui and Muchuan

We detected 10 pupal cells with *C. buqueti* adults or larvae in each area. The reason for the small sample size is that few pupal cells were found in Chishui. Eight adult and larvae died or decayed in the Chishui pupal cells (Figs. 1A–C), but only a few adults were found dead in those obtained from Muchuan (2/10) (Fig. 1 D–F).

**Figure 1.**
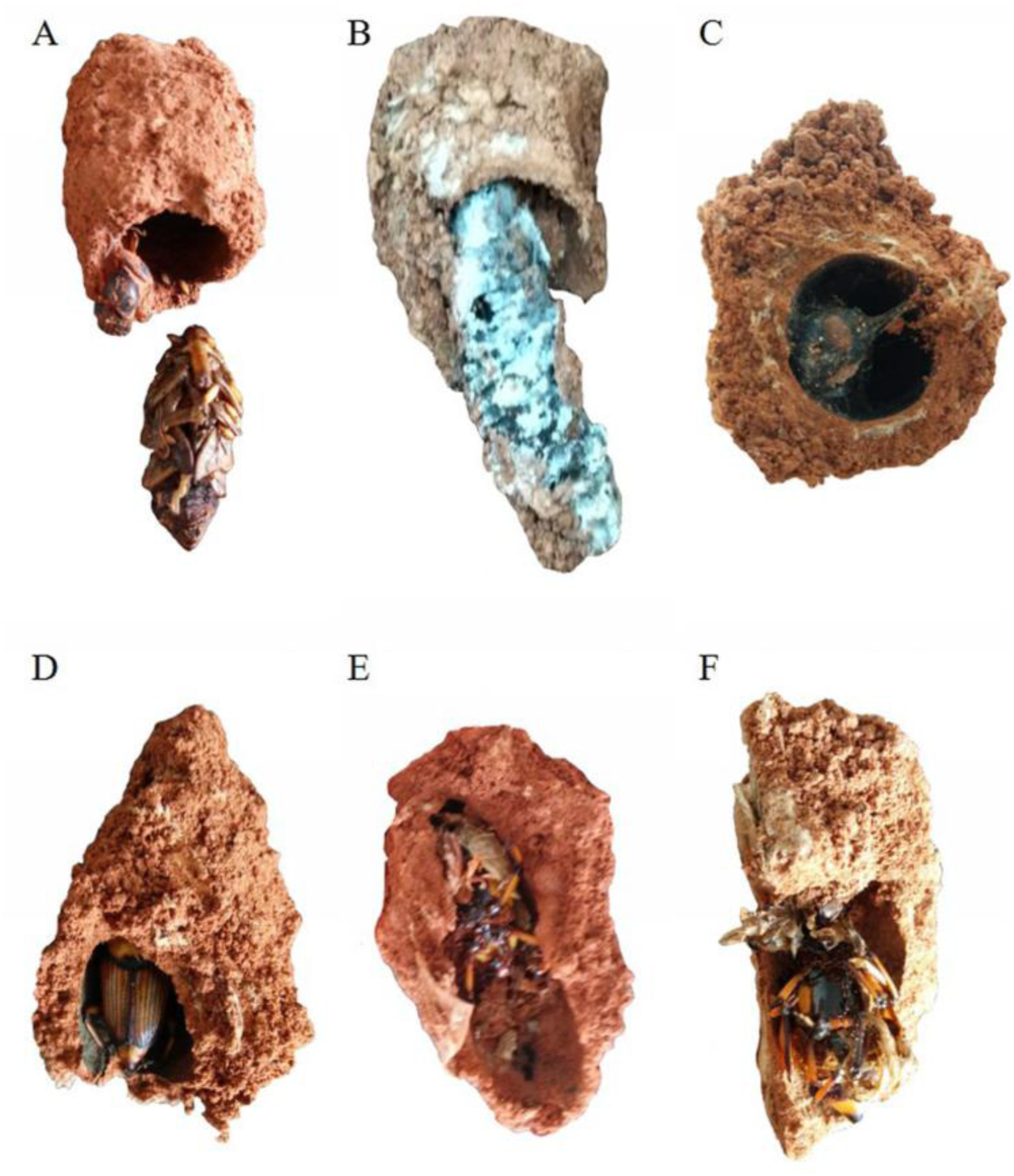
Pupal cells with *Cyrtotrachelus buqueti* adults or larvae in Chishui, Guizhou Province and Muchuan, Sichuan Province. Pupal cells with *C. buqueti* (A–C) adults or larvae from Chishui and (D–F) adults from Muchuan.

The quantitative distribution of *C. buqueti* in Muchuan was much higher than in Chishui. We also found several differences in the living environments of *C. buqueti* between the two regions. Chishui had higher heavy metal concentrations in the soil and pupal cells than Muchuan, with metals such as Hg and Se (Fig. 2A & B). The pH of the soil and pupal cells was higher in Muchuan than in Chishui (Fig. 2C). Moreover, the annual mean temperature in Chishui was 18.1 °C and could reach 28 °C in July. By contrast, the temperature in Muchuan was more than 1 °C lower, averaging at 17.0 °C (Fig. 2D).

**Figure 2.**
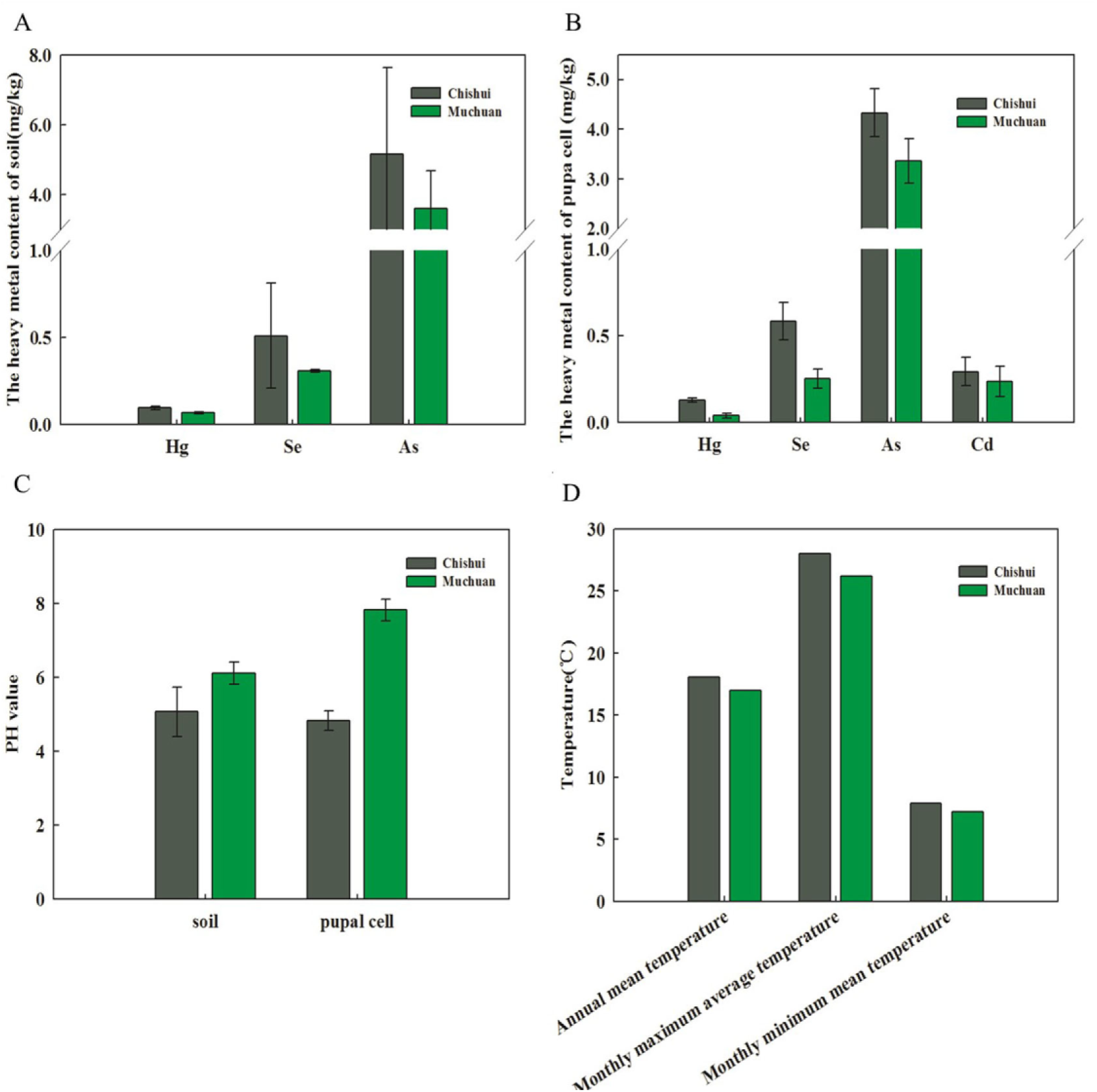
Living environment of *C. buqueti*. Heavy metal content in (A) soil (Se, Hg, and As) and (B) pupal cells (Se, Cd, Hg, and As); (C) pH levels of soil and pupal cells; (D) Temperatures in Muchuan and Chishui, including annual mean temperatures, monthly maximum average temperatures, and monthly minimum mean temperatures.

### 3.2 Sequencing, de novo assembly, and functional annotations

Six sequencing libraries from the two distinct areas were prepared and sequenced with the Illumina HiSeq platform to investigate the population size variation of *C. buqueti*. A total of 52 million short reads were generated from the six libraries, with 44 million high-quality (Q ≥ 20) 125-bp reads, which could be subdivided into 29,406 UniGenes with an average length of 747 bp (Table 2). The lengths of the assembled UniGenes primarily ranged from 201 to 20,229 bp, with a mean length of approximately 748 bp (Fig. 3A). Additionally, a BLAST comparison with the Uniprot protein database revealed that up to 15,641 UniGenes in the *C. Buqueti* transcriptome contained putative homologs in other databases (Fig. 3B). Among the 15,641 UniGenes, 15,703 were successfully annotated by Gene Ontology (GO) assignments and classified into three functional categories, namely, molecular function, biological process, and cellular component (Fig. 3D). The matched unique sequences assigned to molecular function were clustered into 22 classification bins with the largest subcategory being binding (62 UniGenes) and the second largest subcategory being catalytic activity (46 UniGenes). The unique sequences were classified into 23 bins with the most abundant comprising transcripts involved in the cellular process (65 UniGenes), metabolism (54 UniGenes), and single-organism process (54 unigenes) categories. The unique sequences in cellular components were divided into 20 classifications with the most abundant being cell part (75 UniGenes) and organelle (37 UniGenes) (Fig. 3D).

**Table 2.** Summary statistics from Illumina sequencing of the *C. buqueti* transcriptome.

| Sequencing |  |
| --- | --- |
| Total number of reads | 52,439,226 |
| Number of clean reads | 44,036,144 |
| Number of contigs | 97,721 |
| Total length of contigs (bp) | 73,064,686 |
| Contig N50 | 1,588 |
| Average length of contigs (bp) | 330 |
| Number of primary unigenes | 73,210 |
| Number of final unigenes | 29,406 |
| Total length of final unigenes (bp) | 231,348,394 |
| Average length of final unigene (bp) | 1245.5740 |
| GC percentage (%) | 38.4200 |

**Figure 3.**
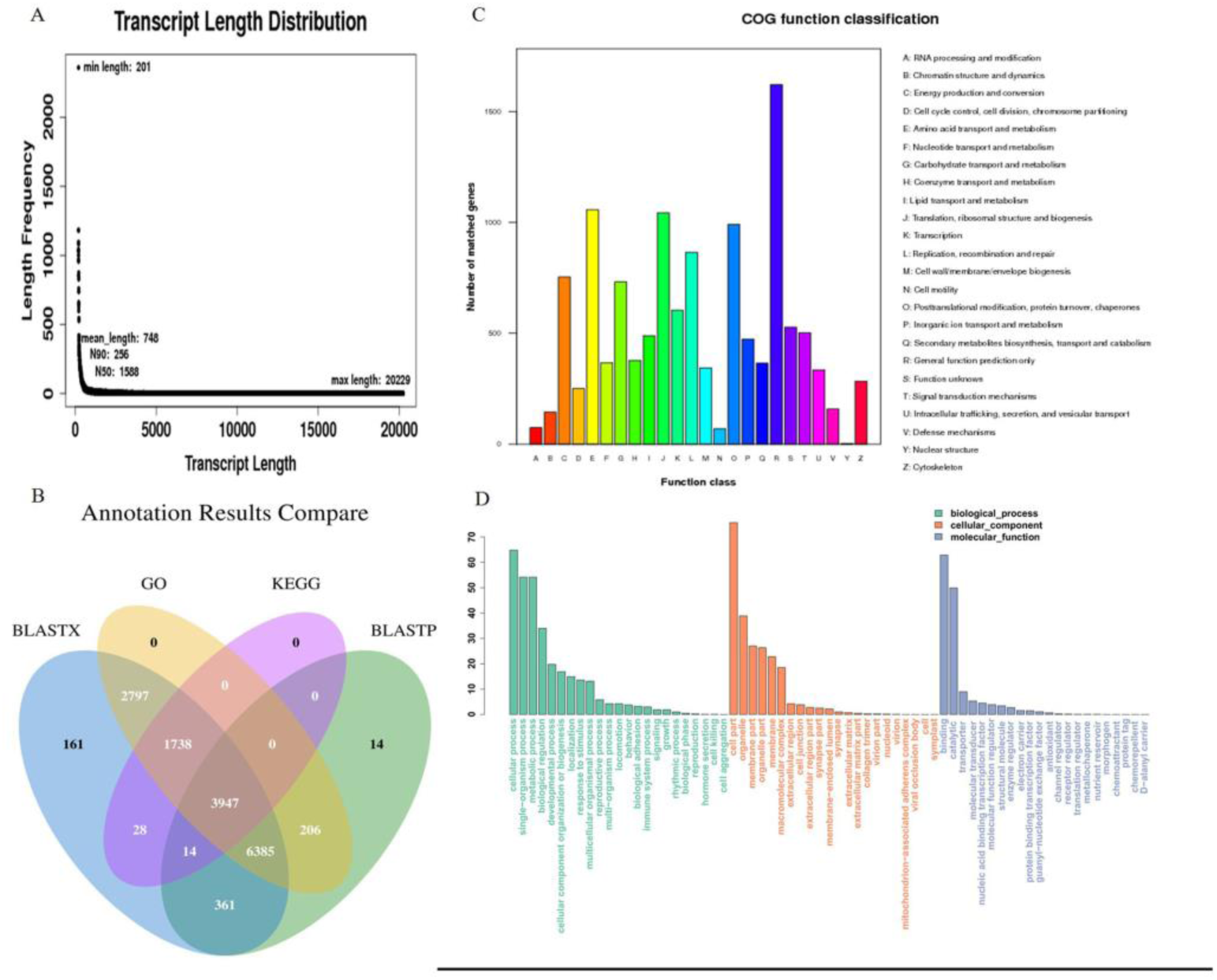
Functional annotations of UniGenes. (A) Length distribution of UniGenes. A total of 15,641 UniGeneswere assembled. (B) Venn Diagram: summary of annotation results. (C) Clusters of Orthologous Groups functional categories in the *C. buqueti* transcriptome. A total of 15,071 UniGenes were subcategorized into 24 categories. The y-axis represented the number of UniGenes, and the x-axis represented categories. (D) Gene Ontology functional categories in the *C. buqueti* transcriptome, including molecular function, cellular component, and biological process.

The matched unique sequences were divided into 24 categories using Clusters of Orthologous Groups (COGs) of proteins (Fig. 3C). The dominant category included general functional prediction, amino acid transport and metabolism, translation, ribosomal structure and biogenesis, post-translational modification, protein turnover, and chaperones (Fig. 3D).

However, the effect of this diverse gene expression on the decreased population density of *C. buqueti* in Chishui remains unknown. First, we performed a hierarchical clustering of the six samples using the Euclidean distance method associated with complete linkage. C2 and C4 were close to each other, similar to D3 and D4 (Fig. 4A), suggesting the use of C2, C3, and C4 or D2, D3, and D4 as three biological repeats. We summarized the expression level of each gene with HT-seq by reads per kilobase million mapped reads (RPKM). The two groups were compared using DEGseq^127^ to identify differentially expressed genes (DEGs). We found 348 DEGs by using a false discovery rate (FDR), 0.05, and a fold change of 2 as significance cutoffs. Among these DEGS, 77 genes were significantly upregulated and 271 were significantly downregulated in the D-group, compared with the C-group (Fig. 4C).

**Figure 4.**
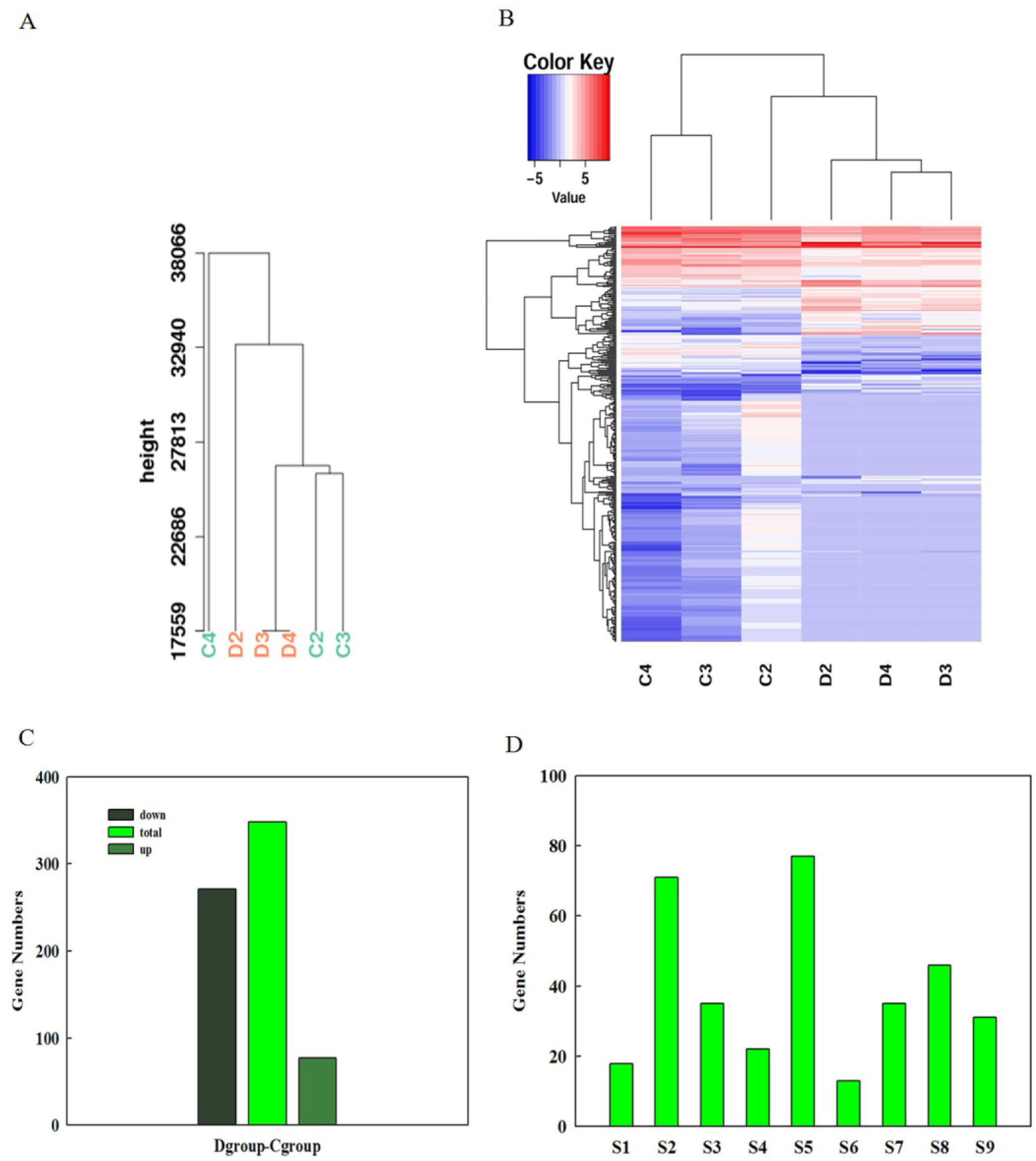
Overview of serial analysis of DEGs identified by pairwise comparisons of the six *C. buqueti* transcriptomes, namely C2, C3, C4, D2, D3, and D4. (A) Cluster of all samples. (B) Heatmap of DEGs across the six *C. buqueti* transcriptomes. The expression values of six libraries are presented as RPKM normalized log 2 transformed counts. Red and blue colors indicate upregulated and downregulated transcripts, respectively. The nine main clusters are shown. (D) Counting the number of each cluster.

We also performed hierarchical clustering of all DEGs by using the R Project for Statistical Computing (Fig. 4B). Nine clusters were plotted with expression patterns (Fig. 4D). The S1 cluster included 18 upregulated genes from both samples, and the expression levels of these genes were similar in both the C- and D-groups. Thirty-two of these genes in the S2 cluster were upregulated in both groups, and the expression levels in the C-group were higher than those in the D-group, and 39 genes had opposite expression patterns. The 35 genes in S3 were downregulated in the D-group but showed no significant changes in the C-group. Interestingly, genes in the S4 cluster had expression patterns contrary to those in S3. The genes in cluster S5 possessed the most DEGs, but gene expression was not significantly different between the two groups. Notably, the gene expression pattern in C4 was distinct from the others. The S6 cluster was composed of 13 genes, the expression of which was downregulated in the C-group compared with the D-group. Genes in the S7–S9 clusters were downregulated in both groups (Fig. 4D).

### 3.3 Functional classification of DEGs between the two groups

We used GO assignments to classify the functions of DEGs in pairwise comparisons of cDNA libraries between the different groups (Fig. 5). Comparative results of the C and D groups showed that many GO terms were significantly enriched in the three categories. The translation and cellular protein metabolic process pathways were significantly enriched in the biological process category (P < 0.05) (Fig. 5A). The ribosome, non-membrane-bounded organelle, intracellular non-membrane-bounded organelle, and ribonucleoprotein component GO terms were significantly enriched in the cellular component category (Fig. 5C). The structural molecule activity and structural constituents of ribosomes terms were significantly enriched in the molecular function category (Fig. 5B).

**Figure 5.**
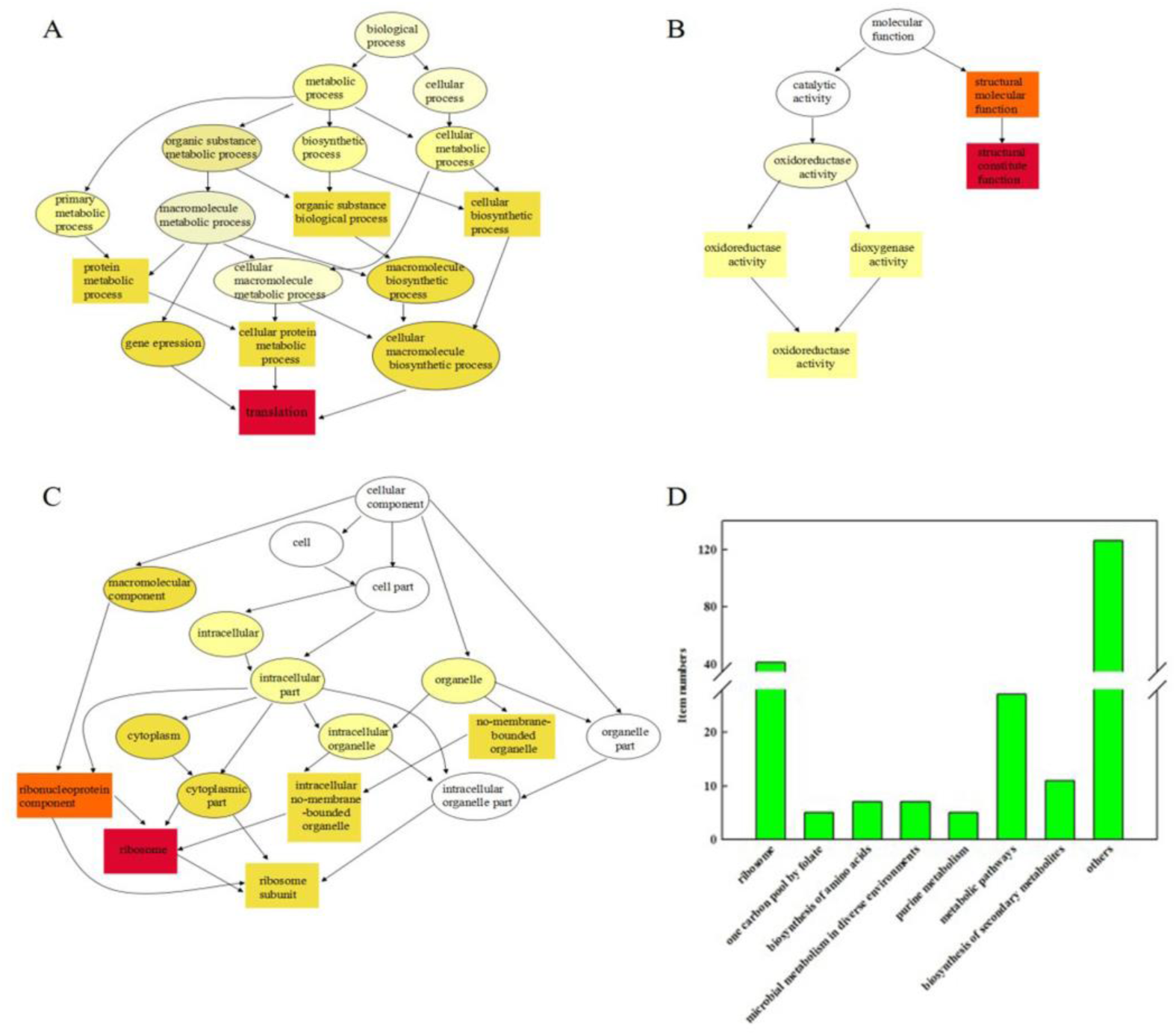
Functional classification of DEGs between the two groups. Directed acyclic graphs of (A) biological process, (B) molecular function, and (C) cellular component categories. (D) Main KEGG Orthology (KO) classifications of DEGs. A total of 348 DEGs were mapped to eight categories; a: ribosome; b: one-carbon pool by folate; c: biosynthesis of amino acids; d: microbial metabolism in diverse environments; e: purine metabolism; f: metabolic pathways; g: biosynthesis of secondary metabolites; and h: others.

We then performed a KEGG analysis of the DEGs. Most of the genes were assigned to the ribosome, one-carbon pool by folate, biosynthesis of amino acids, microbial metabolism in diverse environments, purine metabolism, metabolic pathways, and biosynthesis of secondary metabolite processes categories (Fig. 5D).

### 3.4 Candidate genes involved in one-carbon metabolism, purine synthesis, and folate metabolism

Genes with expressions that were highly correlated with one-carbon pool by folate are potential candidates involved in *C. buqueti* reproduction, growth, and development. We suggested five candidate genes, namely, *AIRS* (c14771_g1_i2), *GARS* (c14771_g1_i1), *SHMT1* (c3484_g1_i2), *SHMT2* (c3484_g1_i1), and *THFD* (c28372_g1_i4), for one-carbon pool by folate. The expression of these genes in the C group significantly differed from their expression in in the D group. The expression levels of candidate genes encoding for *AIRS* and *GARS*, which are key enzymes in IMP synthesis (Ulrich et al., 2003) (Supplementary Fig. 1), were downregulated in the D group (Fig. 6). This suggests that IMP synthesis would be disordered in the D group. Moreover, *SHMT1* and *SHMT2*, which are serine hydroxymethyltransferases and are involved in one-carbon metabolism (Supplementary Fig 1), also had lower expression in the D group than in the C group (Fig. 6). *MTHFR* in the D group, which participates in folate metabolism (Supplementary Fig. 1), was significantly downregulated by twofold (log 2) compared with the C group (Fig. 6). Thus, folate metabolism might be destroyed in the D group. Functional analysis of these candidate genes may be beneficial for determining the molecular mechanisms underlying the disunity of *C. buqueti* quantitative distribution.

**Figure 6.**
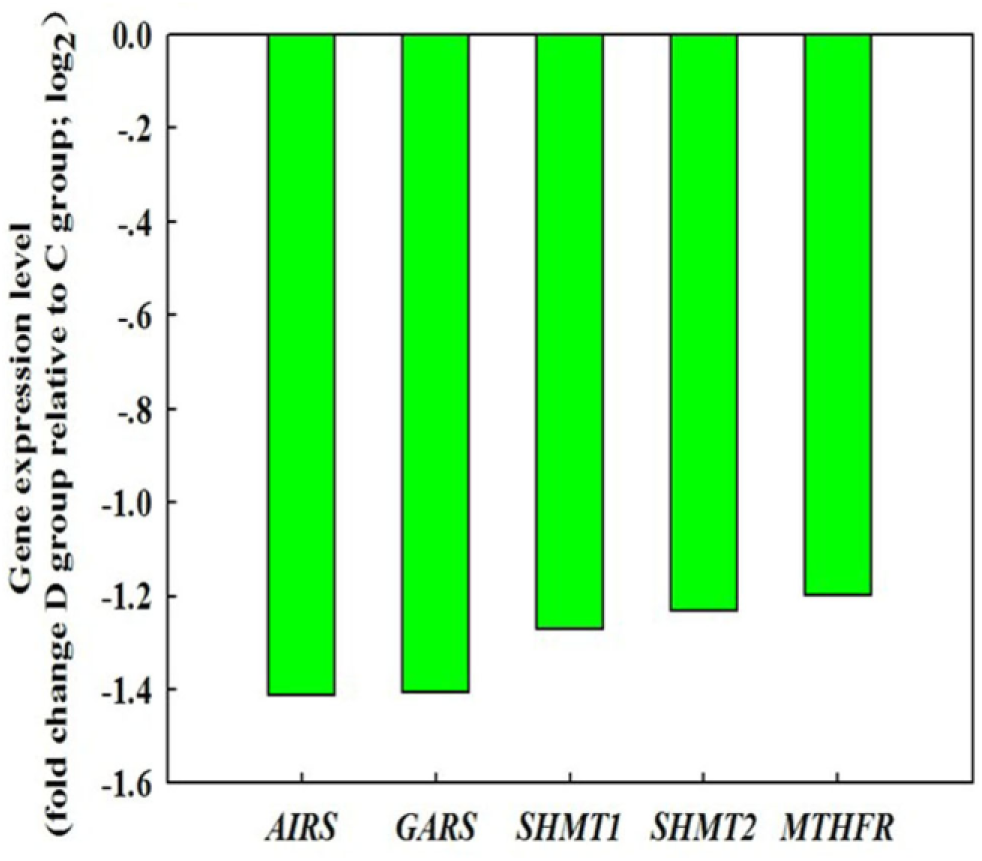
Gene expression levels of *AIRS* (c14771_g1_i2), *GARS* (c14771_g1_i1), *SHMT1* (c3484_g1_i2), *SHMT2* (c3484_g1_i1), and *THFD* (c28372_g1_i4). Twofold (log 2) change of D group relative to C group

### 3.5 Genes related to retinal determination gene network (RDGN)

Genes involved in RDGN have been reported to also be involved in eye development. The genes related to RDGN belong to the PAX6, eyes absent (EYA), Sine oculis(SO), and dachshund (DAC) families (Pappu and Mardon, 2002). In this study, we found 10 PAX6 homologous genes, 10 EYA genes, three SO genes, and four DACH genes (Supplementary Table 1). The expression of these genes in the C group was not significantly different from that in the D group (Supplementary Fig. 2). However, analyzing these RDGN genes may be useful for the determination of the molecular mechanisms for genetic engineering or the marker-assisted selection of *C. buqueti* feeding, which could alter the quantitative distribution of *C. buqueti*.

### 3.6 DEGs involved in the ribosome category

The molecular mechanism involved in the decreased population numbers in Chishui was the downregulation of ribosomal protein genes, resulting in decreased ribosome numbers, and enlarged ribosome size (Korostelev et al., 2006, Zhang et al., 2014). In our study, we found that 41 downregulated ribosomal protein genes were significantly enriched in the ribosome category of the KEGG pathway analysis (Figs. 7A and B).

**Figure 7.**
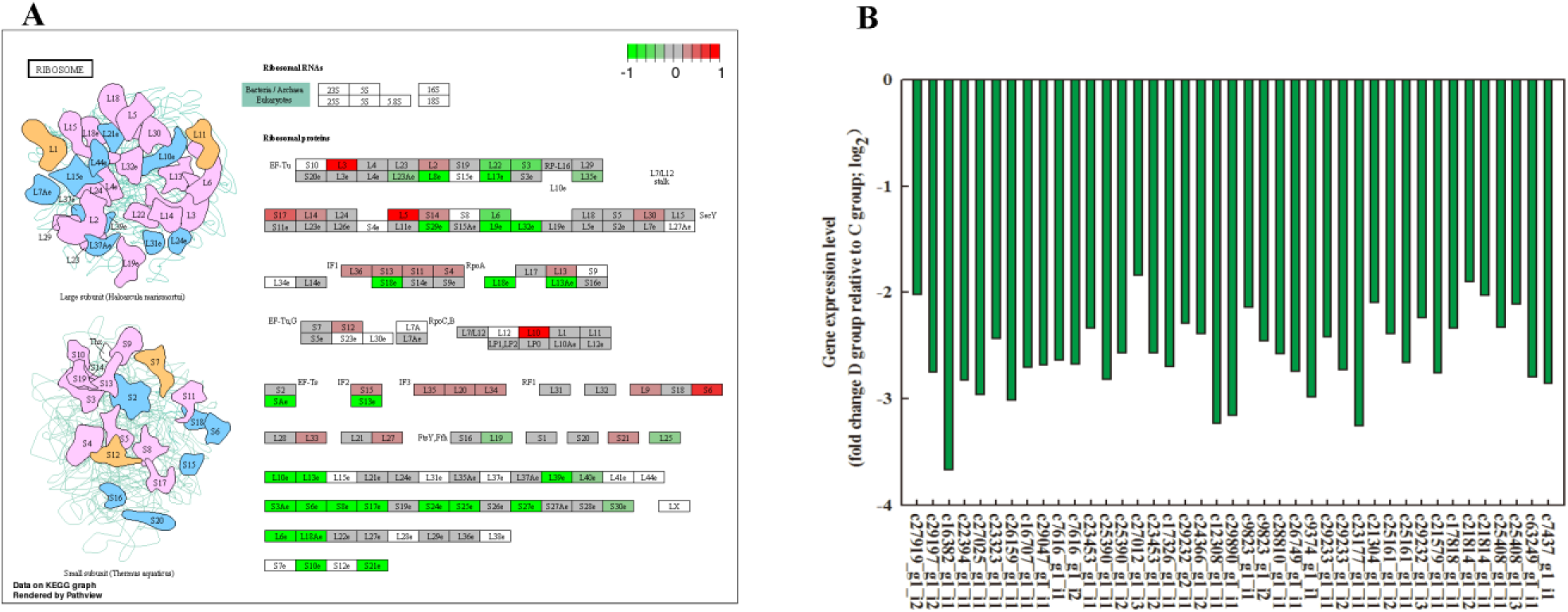
Gene expression levels of ribosomal protein genes. (A) Data on the KEGG graph rendered by pathview. (B) Gene expression level of ribosome protein genes, including 41 genes: c27919_g1_i2, c29197_g1_i2, c16382_g1_i1, c22394_g1_i1, c27025_g1_i1, c23323_g1_i1, c26159_g1_i1, c16707_g1_i1, c29047_g1_i1, c7616_g1_i1, c7616_g1_i2, c23453_g1_i1, c25390_g1_i1, c25390_g1_i2, c27012_g1_i3, c23453_g1_i2, c17326_g1_i1, c29232_g2_i1, c24366_g1_i2, c12308_g1_i1, c29890_g1_i1, c9823_g1_i1, c9823_g1_i2, c28810_g1_i1, c26749_g1_i1, c9374_g1_i1, c29233_g1_i1, c29233_g1_i2, c23177_g1_i1, c21304_g1_i1, c25161_g1_i2, c25161_g1_i1, c29232_g1_i3, c21579_g1_i1, c17818_g1_i1, c21814_g1_i2, c21814_g1_i1, c25408_g1_i1, c25408_g1_i3, c63249_g1_i1, and c7437_g1_i1. Twofold (log 2) change D group relative to C group

### 3.7 Heat shock protein (HSP) relevant gene

Temperature gradients associated with global warming threaten the survival of wild animals, but the expression of HSPs can improve tolerance to heat shock. Five putative homologs of the HSP gene, namely, *HSP1* (c30948_g1_i1), *HSP2* (c30174_g1_i1), *HSP3* (c24612_g1_i2), *HSP4* (c24612_g1_i1), and *HSP5* (c28234_g1_i1), were identified in the transcriptomic data. These genes exhibited lower expression levels in the D group than that in the C group (Fig. 8).

**Figure 8.**
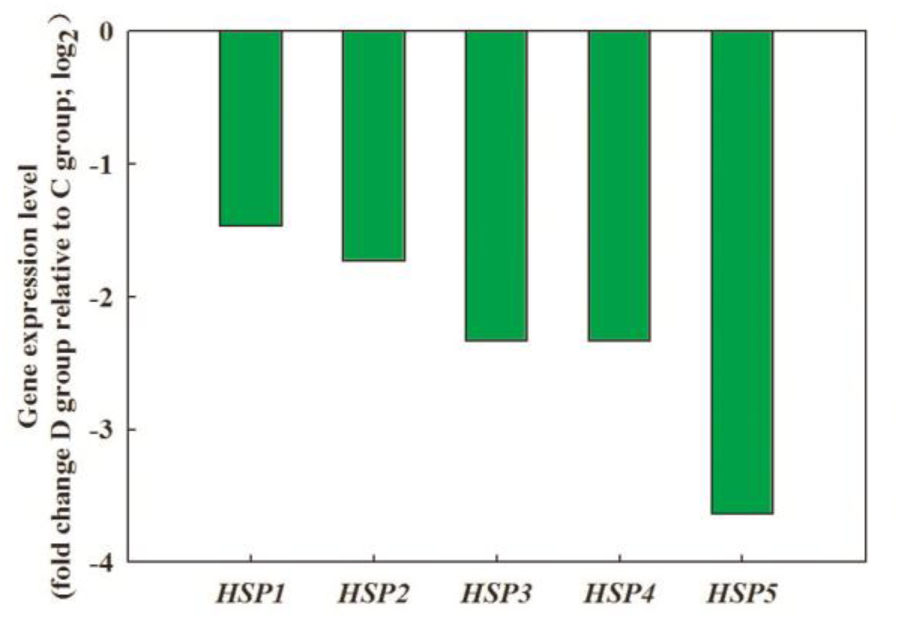
Gene expression level of five HSPs. Five genes, *HSP1* (c30948_g1_i1), *HSP2* (c30174_g1_i1), *HSP3* (c30174_g1_i1), *HSP4* (c30174_g1_i1), and *HSP5* (c30174_g1_i1). Twofold (log 2) change in D group relative to C group.

### 3.8 RT-qPCR validation

The transcriptional regulation revealed by RNA-seq was confirmed in three independent biological experiments using RT-qPCR. A total of 20 genes were selected to design gene-specific primers; 10 were involved in the ribosome pathway, five in the one carbon pool by folate pathway, and five coded HSP (Table 1). The RT-qPCR results for all genes were tested statistically and most genes showed significantly different expression levels (P=0.05, Fig. 9). Moreover, 20 genes showed significant correlations (P=0.05) between the RT-qPCR data and the RNA-seq results, which indicated good reproducibility between the transcript abundance assayed by RNA-seq and the expression profile revealed by RT-qPCR (Fig. 9).

**Figure 9.**
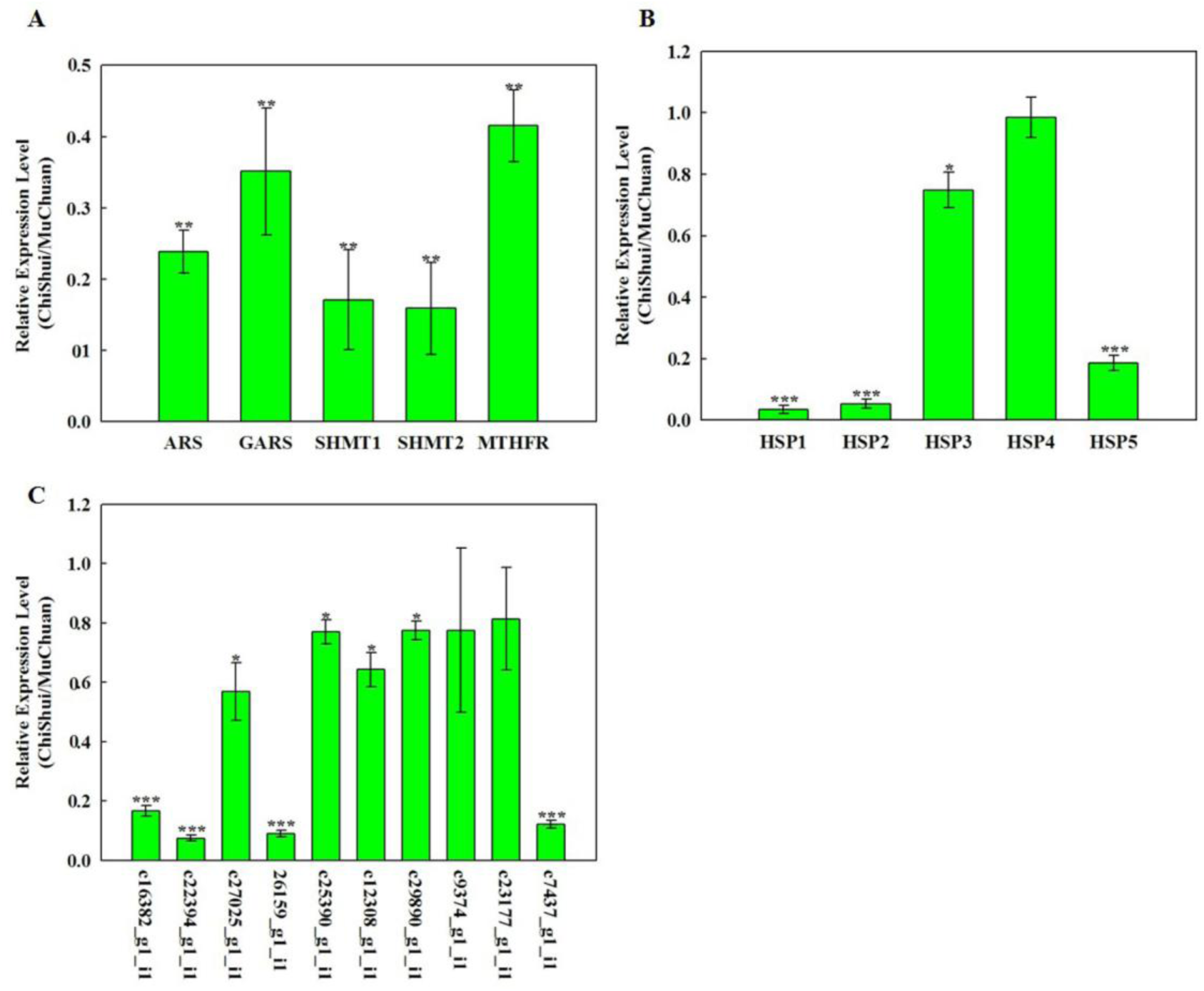
Real-time quantitative RT-qPCR confirmation of 20 candidate genes. The left y-axis indicates relative gene expression levels determined by RT-qPCR. Relative gene expressions were normalized by comparison with the expression of lotus β-actin (c28453_g1_i3), and were then analyzed using the 2^−ΔΔC^T method. The expression values were adjusted by setting the expression of MuChuan samples to be 1 per gene. All RT-qPCRs for each gene used three biological replicates, with three technical replicates per experiment; the error bars indicate SE. Different “*”(*-***) indicates the significant difference at P=0.05. (A) The expression levels of candidate genes involved in one-carbon metabolism, purine synthesis, and folate metabolism. (B) The expression levels of HSPs. (C) The expression levels of candidate genes involved in the ribosome category.

## 4. Discussion

We identified and studied two separate regions with different population sizes of *C. buqueti.* Few were observed in Chishui in Guizhou province, when compared to the abundant population in Muchuan in Sichuan province. The reduced *C. buqueti* numbers are the result of complex factors and are dependent on diverse gene expression controlled by environmental factors. We investigated pupal cells with *C. buqueti* adults or larvae. The results showed that the adults and larvae died in the pupal cells in Chishui, but only a few adults were found dead inside the pupal cells in Muchuan, as predicted. The potential reasons might be that the living environment altered, inducing the formation of several physical and physiological defects in the insects or causing bamboo shoots to become more toxic to the insect. These characteristics shortened the lives of both larvae and adults and even caused death.

Heat is known to cause endogenous (species-native) proteins to misfold into aggregation-prone species, the toxicity of which is mitigated and reversed by chaperones (De, 1995, Vabulas et al., 2010, Verghese et al., 2012). Newly synthesized proteins are particularly susceptible to heat-induced misfolding and aggregation and are apparently major triggers of the heat-shock response, as well as its main beneficiaries (Vabulas et al., 2010). At lower temperatures, more ribosomes are engaged in protein synthesis (Yun et al., 1996). The structures and functions of DNA and RNA are altered at high temperatures, and thus stable heredity can be adversely affected (Liao et al., 2015). We analyzed the temperature patterns in the two regions and found that Chishui had a higher average temperature than Muchuan (Fig. 2D), especially during July and August, with maximum temperatures of over 40°C. Ironically, Nanning, in Guangxi Province, China, which has a higher annual mean temperature than Chishui (data not shown), reported serious damage caused by *C. buqueti.* These results indicate that high temperature only partially, rather than primarily, affects the *C. buqueti* proliferation. In addition, our results showed that Hg, Se, and As existed in higher concentrations in the soil and pupal cells in Chishui, and that the Cd concentration was higher in the pupal cell in Chishui, than was found in Muchuan (Figs. 2A and B). Plants and animals may accumulate heavy metals like Cd, Se, Hg, Cr, and Pb from the soil, which negatively affects protein synthesis, ribosome structure and function, and RNA and DNA synthesis (Ahmad et al., 2014, Wang et al., 2004) A preliminary hypothesis on the mode of action and likely proximal site for organic mercurial inhibition of synaptosome fraction protein synthesis are provided (Cheung and Verity, 1981) in Supplementary Fig 3. Therefore, we assume that a high heavy metal concentration might play a dominant role in reducing the population of *C. buqueti* in Chishui. Moreover, the bioavailability and toxicity of heavy metals is partially dependent on pH. Heavy metals generally became more available to organisms at increased pH levels (Sijm et al., 2000). The results showed that pH levels below 5 in Chishui and up to pH 6 in Muchuan, especially up to pH 7.5 in pupal cells (Fig. 2C), might affect the ribosome-catalyzed peptidyl transfer and the bioavailability and toxicity of heavy metals. In summary, the combined effects of high-temperature, high concentration of heavy metals, and high pH values might combine and adversely affect the diverse metabolism processes of *C. buqueti* by affecting protein, DNA, and RNA synthesis in Chishui samples, which might reduce the quantitative population of *C. buqueti*, and even cause local extinction of the insect population.

The global analysis of transcriptomes could facilitate the identification of systemic gene expression and regulatory mechanisms, in order to successfully analyze the transcriptomes of several species (Xie et al., 2012, Sweetman et al., 2012). In our study, we sequenced and annotated the transcriptome of *C. buqueti* samples from the two study areas to screen for genes responsible for the observed variety of quantitative distribution. A total of 52 million pair-end reads were generated. An average of 94.2% of the data were mapped to the transcriptome (Table 2). Among the 29,406 genes identified, 15,641 were previously predicted in the reference. Moreover, 348 were differentially expressed between the insects in the two different areas, with 77 upregulated and 271 downregulated UniGenes. The functional analysis identified that these genes were significantly enriched in ribosome constitution and metabolic pathways. Candidate genes included 41 involved in ribosome constitution, five in pathway one-carbon pool by folate, and five HSP genes. A comparative transcriptomic analysis detected several DEGs and potential candidate genes. Adding to the currently available expressed sequences would help further the comprehensive understanding of the transcription profiles of the reduced quantitative distribution *C. buqueti*. Transcriptome-wide gene expression profiles were compared between the libraries to identify corresponding genes associated with the variety of *C. buqueti* distribution. DEG expression was remarkably downregulated, and only 77 genes were upregulated in the D group that were not upregulated in the C group (Fig. 4C). These variations may be consistent with the morphological and physiological changes in *C. buqueti*, which were probably due to changes in the living environment of this species. Moreover, nine clusters were plotted with expression patterns that indicated that certain genes could be responsible for the reduced distribution. The GO enrichment analysis revealed that translation, ribosome, and structural constituents of ribosomes were overrepresented terms for DEGs (Figs. 5A–C). These may contribute to the regulation of ribosome protein gene expression, thus affecting ribosome quantity and enlarging ribosome size. KEGG analysis also revealed that most genes were assigned to the ribosome, one-carbon pool by folate, biosynthesis of amino acids, purine metabolism, metabolic pathways, and biosynthesis of secondary metabolite processes categories (Fig. 5D). Thus, these processes play an important role in ribosome quantity, enlargement of ribosome size, and enhancement of enzyme activity.

Folate is a cofactor that transfers single-carbon units (e.g., CH_3_ and CHO) in numerous reactions. One-carbon metabolism includes nucleotide synthesis (purine ring and thymidine from uridine; Supplementary Fig 4 and 1) (Blatch et al., 2015). Thus, insufficient folate would result in the inadequate production of thymidine and possible misincorporations of uracil in DNA (Blount et al., 1997). Insects also appear to require folate for nucleotide synthesis (Blatch et al., 2015). Folate deficiencies decrease SAM levels and DNA methylation levels, which then disrupt gene regulation in mice (Friso and Choi, 2002). Many insects, such as *Drosophila melanogaster*, have methylated DNA, although mechanisms of epigenetic regulation in some insects, including *D. melanogaster*, appear to be quite different from those observed in mammals (Lyko et al., 2000). Studies on folate in insects have provided conflicting results. Folate has been reported to stimulate growth, to have no effect, or even to be toxic. However, insects cannot use folate directly without transforming into THF. 5, 10-Methylenetetrahydrofolate reductase (MTHFR) reduces folate into available THF, which then goes on to participate in DNA synthesis and amino acid transformation. Downregulated *MTHFR* expression causes the accumulation of homocysteine (Hyc) and disordered biochemical processes, such as cell cycle regulation, DNA replication, and DNA and protein methylation (Christensen et al., 1999). We found the following five genes in this study: *AIRS* and *GARS* were the key enzymes of IMP synthesis; *SHMT1* and *SHMT2* were serine hydroxymethyltransferases and are involved in the one-carbon pathway; and the *MTHFR* gene participates in folate metabolism. These genes were downregulated in the D-group (Fig. 6), which can explain the reduced quantitative distribution of *C. buqueti* in Chishui.

Ribosomes are integral parts of any given cell, playing an important role in linking genotypes to phenotypes by manufacturing the proteome. Ribosome construction is essential to maintain cellular homeostasis and is also the single most expensive metabolic process for a cell (Raska et al., 2004). A dividing mammalian cell requires 10 million ribosomes (Purves, 2000), and a rapidly growing bacterial cell requires as many as 20,000 ribosomes (Berg et al., 2006). Therefore, the downregulation of ribosomal protein genes results in the corresponding reduction of ribosome proteins, thus decreasing the quantity of ribosomes in a cell (Korostelev et al., 2006). The enlargement of ribosomes would induce an alteration of biological ribosomal functions and decrease ribosome biogenesis in cells. A previous study showed that cell growth and proliferation could be derived from ribosome biogenesis (Zhang et al., 2014). Thus, ribosome biogenesis is tightly linked with cell growth and proliferation (Trainor and Merrill, 2014, Raiser et al., 2014, Bolze and Casanova, 2013). Our study found that 32 downregulated ribosome protein genes were significantly enriched in the ribosome category of the KEGG pathway analysis (Figs. 7A and B). Therefore, downregulation of ribosomal protein genes may cause alteration in *C. buqueti* cells, such as reduced cell number, cell size, and quantity of cilia. These alterations may have induced an alteration in the quantity of *C. buqueti* in Chishui. However, further studies are needed to elucidate the mechanisms underlying the downregulation of ribosome protein genes and the enlarged size of the ribosomes.

HSPs function as molecular chaperones, and are key in the maintenance of proper protein folding and overall proteostasis, preventing and reversing deleterious protein misfolding (Mayer and Bukau, 2005) (Supplementary Table 2) . Hsp70 assumes a critical role in helping refold aggregated or misfolded proteins, as well as in the folding of newly synthesized proteins, among other functions (Mayer and Bukau, 2005, Höhfeld et al., 2001) (Supplementary Fig 5). It is a crucial component of protein quality control systems, and specialized forms interact with a large continuum of substrates (Höhfeld et al., 2001). Interestingly, five HSP genes were downregulated in the D group, which were the homologous genes of Hsp70 in humans (Fig.8). Moreover, Chishui had a higher average temperature than Muchuan (Fig. 2D). Accordingly, the possibility of protein misfolding might increase, and the overall proteostasis was probably threatened because of the downregulation of HSP genes. These characteristics reduced protein quality and quantity in the *C. buqueti* pupal cells in Chishui. Furthermore, lower expression levels of HSP genes caused lower thermotolerance, whereas the higher temperature in Chishui required a higher thermotolerance of *C. buqueti.* Therefore, *C. buqueti* in Muchuan might respond better to extreme weather, especially to heat. These phenomena indicated that downregulation of HSP genes may decrease thermotolerance and probably result in the observed population size differences between the two regions. However, the mechanisms for the downregulation of HSP genes and amounts of *C. buqueti* remain unknown.

## Conclusions

We explored an effective and safe method to further the molecular understanding of *C. buqueti*. Muchuan and Chishui, which had different *C. buqueti* population sizes, were investigated using transcriptome analysis. Several possible environmental factors, namely temperature, heavy metals and pH value, were found to be responsible for the death of *C. buqueti* adults and larvae in the pupal cells in Chishui, due to hindered metabolic processes, such as protein synthesis, DNA synthesis, RNA synthesis, one-carbon metabolism, purine synthesis, and folate metabolism. Furthermore, we analyzed the expression of RDGE and some HSP genes. The data suggest that the expression of RDGN genes was unchanged, but HSP genes were downregulated in the D group. However, this molecular mechanism did not offer a safer or more effective method for *C. buqueti* prevention, but enriched the molecular knowledge of this pest. Further studies on the molecular mechanisms of feeding behavior, signal molecules, and flight, among others, might help to achieve this goal.

## Competing interests

These authors have no conflict of interest to declare

## Funding

Supported by the National Natural Science Foundation of China (31470655)

## Authors’ contributions

Chaobing Luo and Yaojun Yang analyzed and interpreted the data. Anxuan Liu, Wencong Long and Hong Liao performed the experiment, and was a major contributor in writing the manuscript. All authors read and approved the final manuscript

## Acknowledgements

We would like to thank ANOROAD (Beijing Biotechnology Corporation) for its assistance in origin data processing and related bioinformatics analysis. We also thank Xiang Nong and members of the laboratory for suggestions and discussion of this work and manuscript revision.

